# Women show enhanced attractiveness ratings and neural processing of male faces associated with male chemosensory signals

**DOI:** 10.1101/2020.10.23.352559

**Authors:** Isabelle A. G. Klinkenberg, Christian Dobel, Ann-Kathrin Bröckelmann, Franziska Plessow, Clemens Kirschbaum, Pienie Zwitserlood, Markus Junghöfer

## Abstract

There is growing evidence that humans use olfactory chemosensory signals for social communication, but their role in affective associative learning is largely unknown. To examine this, women implicitly learned face-odor associations by pairing different neutral male faces with either a male chemosignal presumably involved in human mating behavior (dissolved Δ4,16-androstadien-3-one, “AND”), a pleasant smell (dissolved vanillin) or the neutral solvent alone. After learning, women rated faces previously paired with AND or vanillin as more attractive than faces paired with solvent, even though they were unable to identify the contingency of face-odor pairings above chance level. On a neurophysiological level, both AND- and vanillin-associated faces evoked stronger magnetoencephalographic correlates of enhanced emotional attention than solvent-associated faces at early (<120 ms) and mid-latency (140-270 ms) processing stages. This study stresses the role of AND as a human chemosignal in implicit social communication and demonstrates its effectiveness in modulating emotional learning.

## 1 Introduction

Human olfaction is falsely believed to be inferior to that of other mammals (McGann, 2017). In fact, humans can detect virtually all volatile chemicals (>1 trillion), and human olfaction by far outperforms other human senses regarding the number of physically different stimuli it can discriminate (Bushdid et al., 2014). Presumably reflecting differences in sensory environments, different species have different sensitivities to different evolutionarily relevant odorants. In fact, for some odors, human olfaction is orders-of-magnitude more sensitive than that of rodents and dogs (Can Güven and Laska, 2012).

Several studies have demonstrated the importance of body odor for human mating behavior (Havlicek et al., 2008; Herz and Cahill, 1997; Herz and Inzlicht, 2002; Lundström and Jones-Gotman, 2009; Pause et al., 2010; Sergeant et al., 2007; Trivers, 1972). As an example, humans seem to be able to detect, on the basis of body odors, whether potential mates have a similar or different immune system, allowing for the selection of mates that might produce offspring with a more diverse immune system (Milinski et al., 2013). Thus, while it is indisputable that body odors influence individual human sexual behavior, the existence of a general pheromonal communication in humans remains hotly debated.

Originally based on observations in insects, pheromones were defined as “substances which are secreted to the outside by an individual and are received by a second individual of the same species, in which they release a specific reaction, for example, a definite behavior or a developmental process” (Karlson and Lüscher, 1959). Pheromones’ exact molecular mixture is known (Wyatt, 2015), leading to classes of pheromones defined by their function (e.g., alarm (Landolt et al., 1999), food trail (Fitzgerald, 2008) or sex pheromones (Raina and Klun, 1984)).

Yet, due to the large behavioral variability in primates, and also on methodological and theoretical grounds, several authors have refuted the mere existence of pheromones, especially in human primates (Doty, 2010, 2003; Mucignat-Caretta, 2014; Wyatt, 2015; Wysocki and Preti, 2004). In mammals, particularly in humans, learning modulates most behaviors, and, consequently, fixed behavioral responses to pheromones seem unlikely, especially for the more complex behavior involved in selecting mates. Indeed, despite extensive research, a body odor compound that meets all the criteria of a pheromone has not yet been observed in humans (Doty, 2010; Wyatt, 2015).

That said, there is considerable evidence of general chemical olfactory communication between mammals, notably in humans (Lübke and Pause, 2015; McGann, 2017; Zhou and Chen, 2008). Instead of the strictly defined term “pheromone”, many researchers prefer the more neutral term “chemosignal”, an olfactory chemosensory cue serving as a signal in human communication. There is mounting evidence that chemosignals perform several social functions in humans, like threat communication, mother-infant bonding, and modulation of human sexual behavior (for reviews, see Lübke and Pause, 2015; Walla, 2008).

One human chemosignal, by some even considered a human pheromone, is the body odor compound Δ4,16-androstadien-3-one (AND), which seems to be relevant for mating behavior in humans (Sobel and Brown, 2001). AND belongs to a class of volatile steroids produced in the axilla, with stronger concentration in male sweat than female sweat (Gower and Ruparelia, 1993; Preti et al., 1987). AND induces alterations of mood, autonomic arousal, psychological state changes such as enhanced sexual arousal, and higher levels of salivary cortisol in women (Bensafi et al., 2004, 2003; Doty, 1986; Jacob et al., 2001; Jacob and McClintock, 2000; Wyart et al., 2007). Moreover, there is evidence that AND influences basic cognitive functions that facilitate complex social and gender judgments (Saxton et al., 2008; Zhou et al., 2014). As certain brain activation has been found to depend on sexual orientation, a series of studies has shown that smelling AND activates the hypothalamus in heterosexual women and homosexual men, but not in homosexual women or heterosexual men (Berglund et al., 2006; Savic et al., 2005, 2001). Remarkably, AND mixed with solvent is neither particularly pleasant nor unpleasant for most people, but a rather neutral odor.

Here, we further investigated whether AND is unconsciously involved in heterosexual women’s attractiveness ratings of men. We examined in a conditioning paradigm whether females, after being presented with an AND-associated male, would later rate this male as more attractive and react with enhanced attention toward him even in absence of AND (when perceived from distance, or on a photo). Such an enhanced attractiveness rating and attention would support the hypothesis that AND, in addition to the direct and immediate effects reported so far, also has the potential to serve as a so-called unconditioned stimulus (US) in associative learning instances, triggering potentially long-lasting behavioral and neural effects (Mueller and Pizzagalli, 2016). Toward this aim, we showed female participants photos of neutral male faces for evaluation, where the photos were presented along with either AND dissolved in propylene glycol or the solvent alone (SOLV), which served as a control condition.

If the experiment shows behavioral and neural correlates of enhanced attention toward AND-associated faces, this would would support our hypothesis, whereas negative findings would leave us with the question of whether AND has no potential as an unconditioned stimulus or whether appetitive olfactory conditioning per se might just not work in our specific laboratory setting (see below), even if the US was consciously perceived as pleasant. Therefore, we introduced a second control condition by pairing a third set of male faces with a consciously perceptible pleasant odor. As no human body odor is reliably consciously perceived as appetitive, we opted for vanillin (VAN) as a non-social odor. VAN is reliably perceived as pleasant (Seubert et al., 2009), and it is frequently used in neuropsychological experiments (Frasnelli et al., 2011; Kettenmann et al., 1996; Sobel et al., 1998) and as an odorant in women’s and men’s fragrances.

As an experimental paradigm, we chose MultiCS conditioning, which has previously been used in several sensory domains (Bröckelmann et al., 2011; Steinberg et al., 2013b, 2012). Classical or evaluative affective conditioning typically uses one or few neutral stimuli (conditioned stimulus, CS) that become associated with an unconditioned stimulus (US) within a few contingent CS-US pairings; thereby, it acquires the power to elicit a conditioned response (CR) (Andreatta and Pauli, 2015; De Houwer et al., 2001; Martin-Soelch et al., 2007). In contrast, MultiCS conditioning involves multiple complex stimuli (e.g., faces or tones) per affective category that acquire emotional meaning within a few associative-learning trials. MultiCS conditioning is especially suitable for recording event-related potentials or magnetic fields (ERPs/EMRFs), because even a few repetitions of each CS provide a large number of trials within experimental conditions. With the 78 different face identities (26 per US odor) used here, we clearly overcharge processing with contingency awareness. Our implicit learning variant resembles typical naturalistic visual-olfactory associations, and it prevents complex interactions of potentially implicit face-AND and explicit face-VAN pairings, which is not within the scope of this study.

To assess the neurophysiological correlates of effective implicit associative affective learning, we recorded magnetoencephalographic (MEG) brain responses evoked by the CS-faces before and after learning. MEG was chosen because it offers a direct and reference-free measure of neural activity with excellent temporal and good spatial resolution.

Neurophysiological changes in response to rather short phases of associative learning are typically reflected by mid-latency difference components in the time range between 120 and 300 ms. The spatio-temporal occurrence of mid-latency conditioning effects resembles ERP difference components, which have been shown for a wide variety of emotional stimuli such as emotional scenes (Schupp et al., 2006), emotional faces (Mavratzakis et al., 2016; Morel et al., 2009; Rellecke et al., 2013; Schupp et al., 2004), emotional words (Kissler et al., 2007), and emotional gestures (Flaisch et al., 2009). This difference component – termed early posterior negativity (EPN) – is assumed to predominantly reflect enhanced motivated attention toward salient stimuli, eventually leading to feedback loop-driven, amplified sensory processing. Its magnetic counterpart, the EPN-m, is characterized by increased ingoing fields (here defined as negativity) over the right hemisphere and increased outgoing fields (here, positivity) over the left hemisphere (i.e., roughly reflecting the 90° right hand rotation of the electric EPN topography) for affective relative to more neutral stimuli.

Based on previous findings of affective associative face learning, we expected to find an EPN-m difference component for AND- and VAN-associated faces but not SOLV-associated faces. As this EPN-m component would mainly reflect an enhanced processing of and emotional attention toward emotionally salient faces relative to neutral faces, we would not expect major qualitative differences between AND- and VAN-associated faces. Quantitative differences may reflect differences in saliency and depend on factors such as ecological validity, differential impact of social vs. non-social US, or differential impact of implicit vs. explicit perception. Thus, it is impossible to predict whether AND or VAN conditioning would result in stronger saliency. As first step, here we test the prediction that both AND and VAN reveal enhanced EPN-m effects compared to SOLV-associated faces.

Moreover, the EPN/EPN-m is often accompanied by early components, even as early as 60 to 90 ms after stimulus onset, with similar topographies and polarities as the EPN/EPN-m (Bröckelmann et al., 2011; Junghöfer et al., 2015; Steinberg et al., 2013b, 2013a, 2012; Stolarova et al., 2006). Therefore, to assess possible early difference components of emotional associative learning, we also analyzed ERMF earlier than 120 ms.

Additionally, to determine whether the effects of implicit associative learning would be strong enough to modulate behavior, we also measures participants’ ratings of CS-face hedonic valence and emotional arousal before and after the learning phase.

## 2 Methods

### 2.1 Participants

Twenty heterosexual females (19-35 years, M = 25.2, SD = 4.06) participated in a within-subject associative-learning experiment. All were German native speakers, right-handed, and not pregnant. Their vision was normal or corrected to normal. Participants had taken no medication that might influence their sense of smell during the previous three months and had not used hormonal contraception. They were non-smokers and had never suffered from anosmia. They signed an informed consent according to the Declaration of Helsinki and were paid for participation. This study was approved by the Ethics Committee of the Department of Psychology and Sports Science at the University of Muenster, Germany. For more detail, please refer to the supplemental information.

### 2.2 Experimental stimuli and apparatus

Seventy-eight neutral male faces were used as CS. They were obtained from various face databases (The Karolinska Directed Emotional Faces (Lundqvist et al., 1998), the NimStim faces (Tottenham et al., 2009), the Feret database (Phillips et al., 2000, 1998) and from our own institute’s database. The hair in the images was cropped, and pictures were converted to grayscale using Adobe Photoshop CS6 (Adobe, San Jose, CA, USA). Propylene glycol (neutral odor; Sigma-Aldrich, St. Louis, Missouri, USA), AND (suggested positive chemosignal, 46 ppm dissolved in propylene glycol; Steraloids Inc., Newport, Rhode Island, USA), and VAN (positive odor, 400 mg dissolved to 10 ml propylene glycol; Sigma-Aldrich, St. Louis, Missouri, USA) were used as US.

A custom-made olfactometric stimulator described in Steinberg et al. (2012) was used for odor presentation. The flexible tubes that oriented the odorated airflow toward the participant’s nostrils were made of the inert material Teflon to prevent chemical reactions between odors and tubes. To assess the ideal moment of odor presentation, participants wore a custom-made respiration belt that measured chest extension during inhalation. Inhalation frequency and amplitude was thus assessed, allowing US odor presentation during inhalation.

### 2.3 Paradigm

During all parts of the experiment, stimulation was provided through Presentation® (Neurobehavioral Systems, California, US).

#### 2.3.1 Behavioral measurements and self-report data collected pre- and post-conditioning

To prevent novelty and reduce order effects, all 78 CS-faces were first presented once without task to the participants. Each CS-face appeared for 1500 ms without inter-stimulus intervals (ISI). Next, participants rated the hedonic valence and emotional arousal of the faces using a computerized version of the self-assessment manikin, once before (pre) and once after (post) the MEG-measurements (SAM-rating; Bradley and Lang, 1994). After the post SAM-rating and after having been informed about the purpose of the experiment and the kind of odors presented as US, participants were asked to recognize the odors. Each odor was randomly presented five times, and participants were asked to identify them as either SOLV, AND or VAN. Participants were also asked to rate the odors’ hedonic valence on a SAM-scale, and the odors’ intensity on a scale ranging from 0 (not detectable) to 100 (maximum intensity). To test participants’ contingency awareness of CS-US pairings, 15 CS-faces that had previously been paired with either AND, VAN or SOLV were presented again. After each CS presentation, participants were asked to indicate with which odor the face had previously been paired. See Fig. 1A for an illustration of the paradigm.

**Figure 1:**
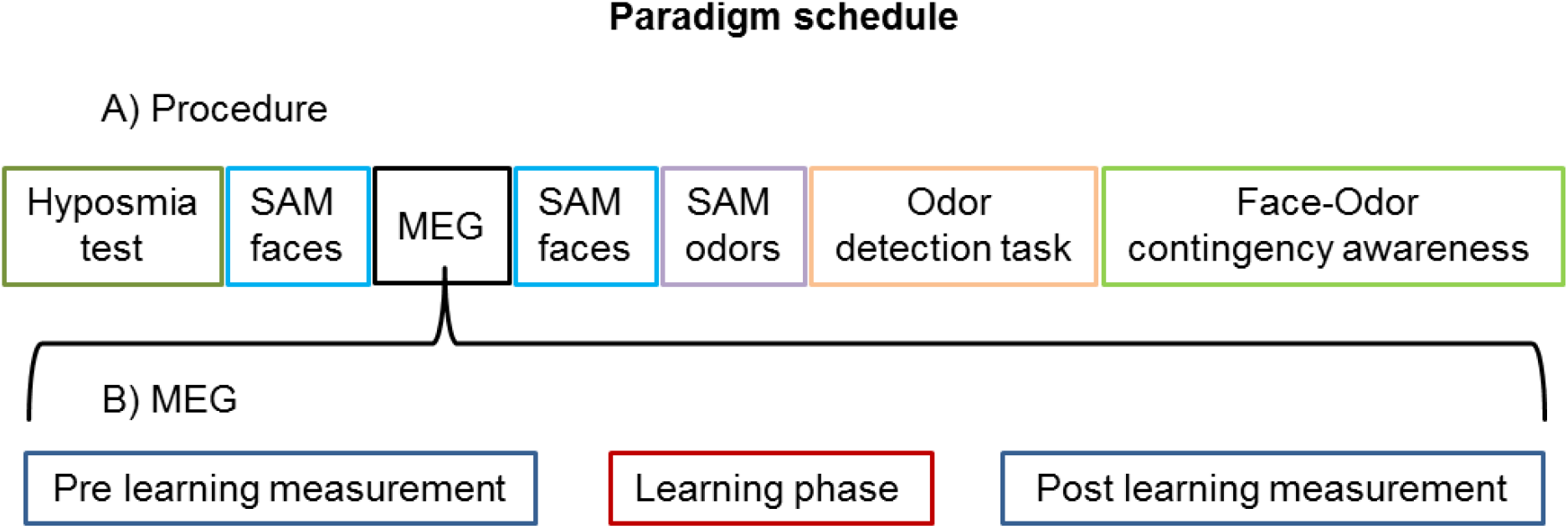
A) Sequence of behavioral, self-report, and MEG measurements and stimulations. B) Description of phases presented during the MEG measurement.

#### 2.3.2 MEG measurements and conditioning

Individual head positions in the MEG were tracked by three electrical landmark coils attached to the two auditory canals and the nasion. A Polhemus 3Space Fasttrack system (Polhemus, Colchester, VT, USA), measuring individual head shape, was used to specify the individual head coordinate system. Participants were placed in the MEG scanner, consisting of a 275-channel MEG whole-head sensor system (Omega 275, CTF, VSM Medtech Ltd., Coquitlam, British Columbia, Canada), with first-order axial SQUID (Super-conducting QUantum Interference Device) gradiometers enabling the measurement of visually evoked magnetic fields (VEMFs).

The MEG part of the experiment was composed of three consecutive phases: a pre-learning, a learning (conditioning), and a post-learning phase (Fig. 1B). Before and after learning, CS-faces were randomly presented for 500 ms (ISI = 750-1500 ms) three times each (3 x 78 faces) in a passive viewing setting. In the learning phase, separate sets of 26 CSs were assigned to each of the three USs: AND, VAN, and SOLV. This assignment was balanced across participants. Each CS-face was paired three times with its assigned US, resulting in overall 234 learning trials. US order was pseudo-randomized, such that the same odor was not presented consecutively. US and CS onset were presented simultaneously at the measured inhalation onset^1^, with a presentation duration of 1000 ms for the US and 2000 ms for the CS^2^. At the beginning of the next inter-trial interval (ITI) – which lasted for a minimum of 1500 ms and at maximum until the next inhalation, and during which a white fixation cross was presented – clean air was projected for 1000 ms in order to neutralize the preceding odor. It is important to note that since odors were randomly presented and the probabilities of odor transitions were controlled, complete eradication of preceding US odors was not necessary. At a height of 15 cm and at a distance of 86 cm between a participant’s nasion and the screen, the face images covered a visual angle of approximately 10° (image border to image border).

## 3 Analysis

### 3.1 Self-report and behavior

Ratings of the odors’ valence scores were *t*-tested against the neutral SAM-value 5. Paired *t*-tests for valence/intensity scores of AND vs. VAN, AND vs. SOLV, and VAN vs. SOLV were calculated, as were linear regressions of odor valence/intensity scores and neural effects. Odor recognition performance was analyzed with the sensitivity measure *d’* (Green and Swets, 1966), computed for the three odors AND, VAN, and SOLV, and tested against 0 with *t*-tests.

To assess face-valence and arousal differences induced by the odor conditioning, pre-conditioning face ratings were subtracted from post-conditioning face ratings. Planned contrasts (AND > SOLV and VAN > SOLV: +1 +1 −2) were conducted to test the hypotheses that both AND- and VAN-paired faces achieve higher valence and arousal ratings in the post-conditioning face ratings^3^. Moreover, paired *t*-tests for AND vs. VAN, AND vs. SOLV, and VAN vs. SOLV were calculated. Due to technical problems, data from one participant had to be excluded from the analysis; therefore, data from 19 participants were included in the Pre-Post SAM-rating analysis of the faces.

For contingency awareness of CS-US pairings, we computed *d’* for AND, VAN, and SOLV faces and *t*-tested each of them against 0. Because of technical issues, data from three participants had to be excluded, resulting in 17 participants in the contingency awareness analysis. Linear regressions of odor valence/intensity scores and neural effects were calculated using R (version i386 3.3.3). All other statistical tests were calculated using SPSS Statistics 22 (IBM, Armonk, New York, US).

### 3.2 MEG

VEMFs induced by the face presentation during the pre-learning and post-learning phases were recorded continuously in a frequency band between 0 and 150 Hz and were digitalized with a sampling rate of 600 Hz. Recordings were offline zero-phase filtered to a bandwidth of 0.1 to 48 Hz (Butterworth second-order high-pass and fourth-order low-pass). Extracted epochs were 800 ms long (ranging from 200 ms before to 600 ms after face onset) and were aligned to stimulus onset. The baseline was adjusted using a 150 ms pre-stimulus interval. For single-trial editing and artifact correction, we applied the method by Junghöfer and colleagues (2000) for statistical control of artifacts in high-density EEG/MEG. This procedure detects individual channel and global artifacts and replaces artifact-contaminated sensors by spline interpolations while computing signal variance across trials as an estimate of stability of the averaged waveform. Subsequently, within each individual, epochs of visually evoked magnetic fields (VEMFs) were averaged depending on US (SOLV, AND, VAN) and phase (pre-learning, post-learning). On average, 72.7 (71.5-73.5) of the total 78 trials (93%) for each experimental condition remained for final averaging. The number of remaining trials did not differ across experimental conditions (F(5,114) = 0.52, p = .77)

Similar to the behavioral data, the averaged MEG data collected before learning (pre-learning phase) were subtracted from the averages of the post-learning phase, and these difference fields were tested via planned contrasts (AND > SOLV and VAN > SOLV: +1 +1 −2). The method described by Maris and Oostenveld (2007) served as a non-parametric statistical testing approach to consider potential deviations from the normal distribution and to correct for multiple comparisons.

To be considered statistically significant, effects had to meet both a sensor-level and a cluster-level criterion. The sensor-level criterion defined that only spatio-temporal data (sensors at given time points) at which the above-planned *F*-contrasts exceeded a critical alpha-level of *p* = .05 were considered for subsequent analysis. To compute the cluster-level criterion, we first identified spatially and temporally adjacent sensors with continuous sensor-level significant activation that formed a spatio-temporal cluster and then calculated the cluster mass, which was the sum of *F*-values within the cluster. Next, Monte Carlo simulations of identical analyses were performed, with 1,000 permutated drawings of experimental conditions. To be included in subsequent analysis, a cluster’s mass had to exceed 99% of the Monte Carlo-simulated cluster masses (maximal cluster mass per drawing), which corresponds to a cluster-level criterion of *p* = .01. To check for effect directions in significant clusters, post hoc ANOVAs were conducted. Differences between single-odor conditions were tested with one-tailed planned contrasts according to our hypotheses (AND > SOLV and VAN > SOLV). Based on our hypotheses and previous results, two time frames were analyzed: an early period of 0 to 120 ms after stimulus onset, and a mid-latent period of 120 to 300 ms after stimulus onset. For the resulting significant clusters, post hoc paired *t*-tests for VEMF amplitudes of AND vs. VAN, AND vs. SOLV, and VAN vs. SOLV were computed. For MEG data preprocessing, artifact rejection and correction, averaging, statistics, and visualization, the MATLAB-based EMEGS software was used (Peyk et al., 2008). Post hoc paired *t*-tests and planned contrasts were performed with SPSS Statistics 22. See supplemental information for the analysis and results of the neural source estimation (Table S1; inverse source modeling).

## 4 Results

### 4.1 Self-report and behavior

The odors AND and SOLV were rated as neutral (AND M = 5, SD = 2.23; SOLV M = 5, SD = 1.41), and ratings did not differ from the neutral SAM value 5 (*t*(19) = 0.00, *p* = 1 for both *t*-tests). VAN, by contrast, was rated as appetitive (M = 6.75, SD = 1.77), as its rating was significantly higher than 5 (*t*(19) = 4,41, *p* < .001) and was also significantly higher than both the AND and SOLV ratings (VAN vs. AND *t*(19) = −3.49, *p* = .002; VAN vs. SOLV *t*(19) = 3.2, *p* = .005; Fig. 2A, left). Odor intensity ratings revealed that AND and VAN were both perceived as more intense than SOLV alone (*t*(19) = 4.58 AND vs. SOLV, *t*(19) = 5.83 VAN vs. SOLV, *p* < .001 for both *t*-tests), whereas the higher intensity ratings for VAN (mean = 70.25; SD = 17.88) than for AND (mean = 64.15, SD = 21.83) were not significant (*t*(19) = −1.29, *p* = .212; Fig. 2A, right).

**Figure 2:**
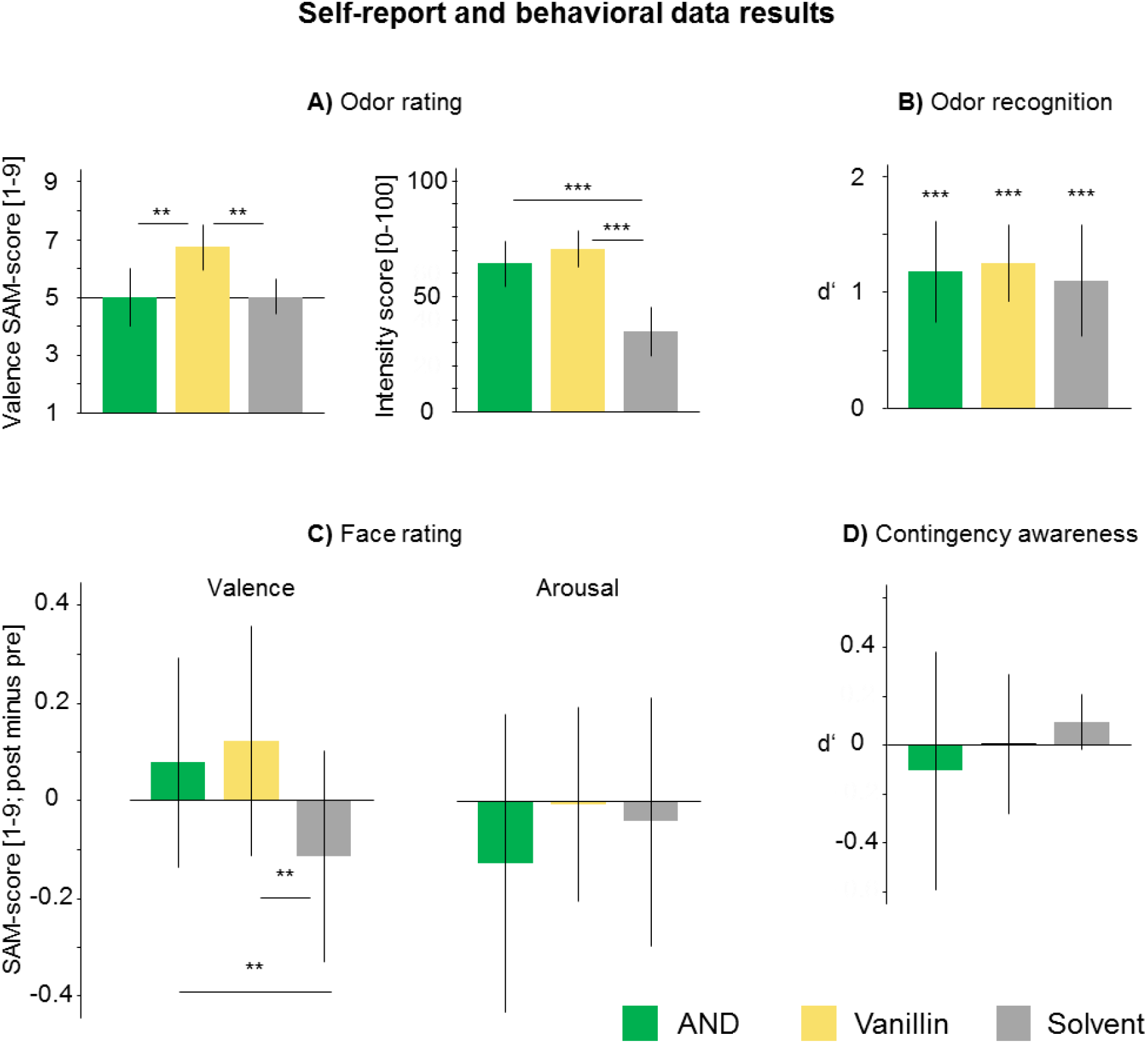
Results of self-report and behavioral data for (A) odor rating (valence SAM-rating left, with the absolute neutral value 5 marked by a horizontal line; intensity rating right), (B) odor recognition task, (C) face SAM-rating (post-rating minus pre-rating), and (D) contingency awareness task. Error bars represent ± 2 SE.

The odor-recognition task showed that *d’* was significantly higher than 0 (AND *t*(19) = 5.35, VAN *t*(19) = 7.44, SOLV *t*(19) = 4.55, *p* < .0001 for all *t*-tests against 0), indicating that participants could identify AND, VAN, and SOLV (Fig. 2B).

For the face-valence ratings (post-minus pre-conditioning), the planned contrast analysis (AND > SOLV and VAN > SOLV: +1 +1 −2) was significant (*F*(1,18) = 11.44, *p* = .003). Post hoc *t*-tests revealed that AND- and VAN-paired faces were rated as more pleasant in the post-conditioning face ratings than SOLV-paired faces (AND vs. SOLV *t*(18) = 2.3, *p* = .034; VAN vs. SOLV *t*(18) = 2.32, *p* = .032), while valence ratings of AND- and VAN-associated faces did not differ (AND vs. VAN *t*(18) = −.32, *p* = .75; Fig. 2C, left). The planned contrast analysis for the arousal ratings and all post hoc comparisons were not significant (F <1 and all *t*s <1; Fig. 2C, right).

Supporting the hypothesis that participants lacked contingency awareness of CS-US pairings, *d’* values did not differ from 0 (AND faces *t*(16) = −.44, *p* = 0.668; VAN faces *t*(16) = .02, *p* = 0.988, SOLV faces *t*(16) = 1.61, *p* = 0.127; Fig. 2D).

### 4.2 Visually evoked magnetic fields (VEMFs)

#### 4.2.1 Early time interval

In the early time frame (0-120 ms), three widely distributed spatio-temporal clusters achieved significance (critical cluster mass (CCM): 1301; Fig. 3A and Fig. 4). The first cluster was located over the left frontal hemisphere and lasted from 27 to 118 ms (cluster mass (CM): 4269.8; Fig. 3A, left; Fig. 4A). Increased outgoing fields (positivity) for both AND- and VAN-associated faces, compared to SOLV-associated faces, were observed in this cluster (AND vs. SOLV *t*(19) = 4.3, *p* < .0001; VAN vs. SOLV *t*(19) = 2.83, *p* = .011), while magnetic fields evoked by AND- and VAN-associated faces did not differ (AND vs. VAN *t*(19) = .32, *p* = .702). Two additional spatio-temporal clusters emerged over the right hemisphere, one more posterior, which lasted from 15 to 62 ms (CM: 1632.9; Fig. 3A, center; Fig. 4B), the other more frontal, lasting from 57 to 97 ms (CM: 2694.7; Fig. 3A, right; Fig. 4C). In both right hemispheric clusters, increased ingoing fields (negativities) were observed for both AND- and VAN-associated faces, compared to SOLV-associated faces (right posterior cluster AND vs. SOLV *t*(19) = −3.62, *p* = .002; VAN vs. SOLV *t*(19) = −2.93, *p* = .009; right frontal cluster AND vs. SOLV *t*(19) = −3.48, *p* = .002; VAN vs. SOLV *t*(19) = −4.32, *p* < .0001), while fields evoked by AND- and VAN-associated faces did not differ (right posterior cluster AND vs. VAN *t*(19) = −.53, *p* = .602; right frontal cluster AND vs. VAN *t*(19) = .24, *p* = .813).

**Figure 3:**
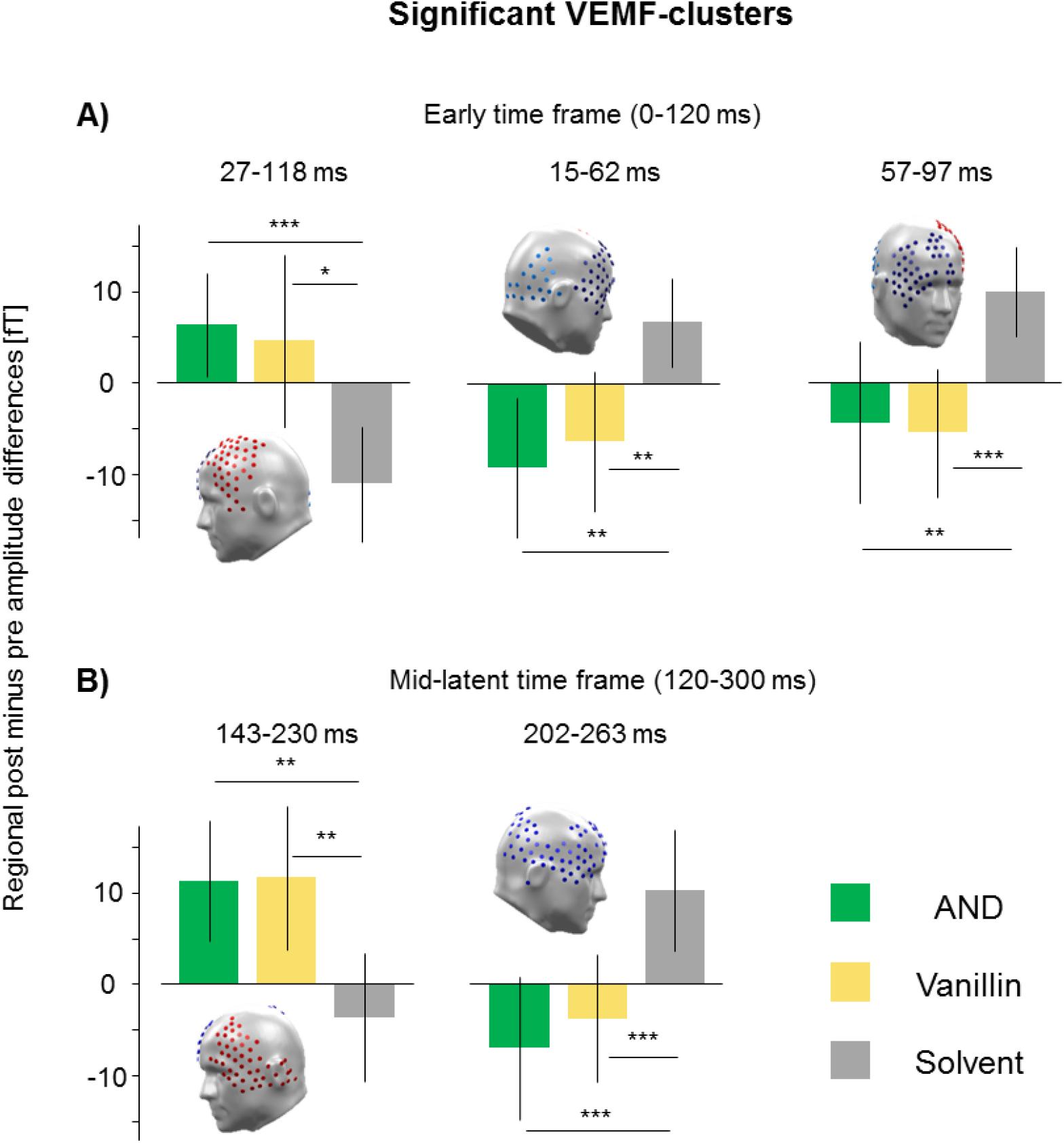
VEMF clusters within (A) the early (0-120 ms) and (B) the mid-latency (120-300 ms) EPN-m time interval. Depicted heads show plotted significant clusters, with red dots representing sensors over the left hemisphere, and blue dots representing sensors over the right hemisphere. Error bars represent ± 2 SE.

**Figure 4:**
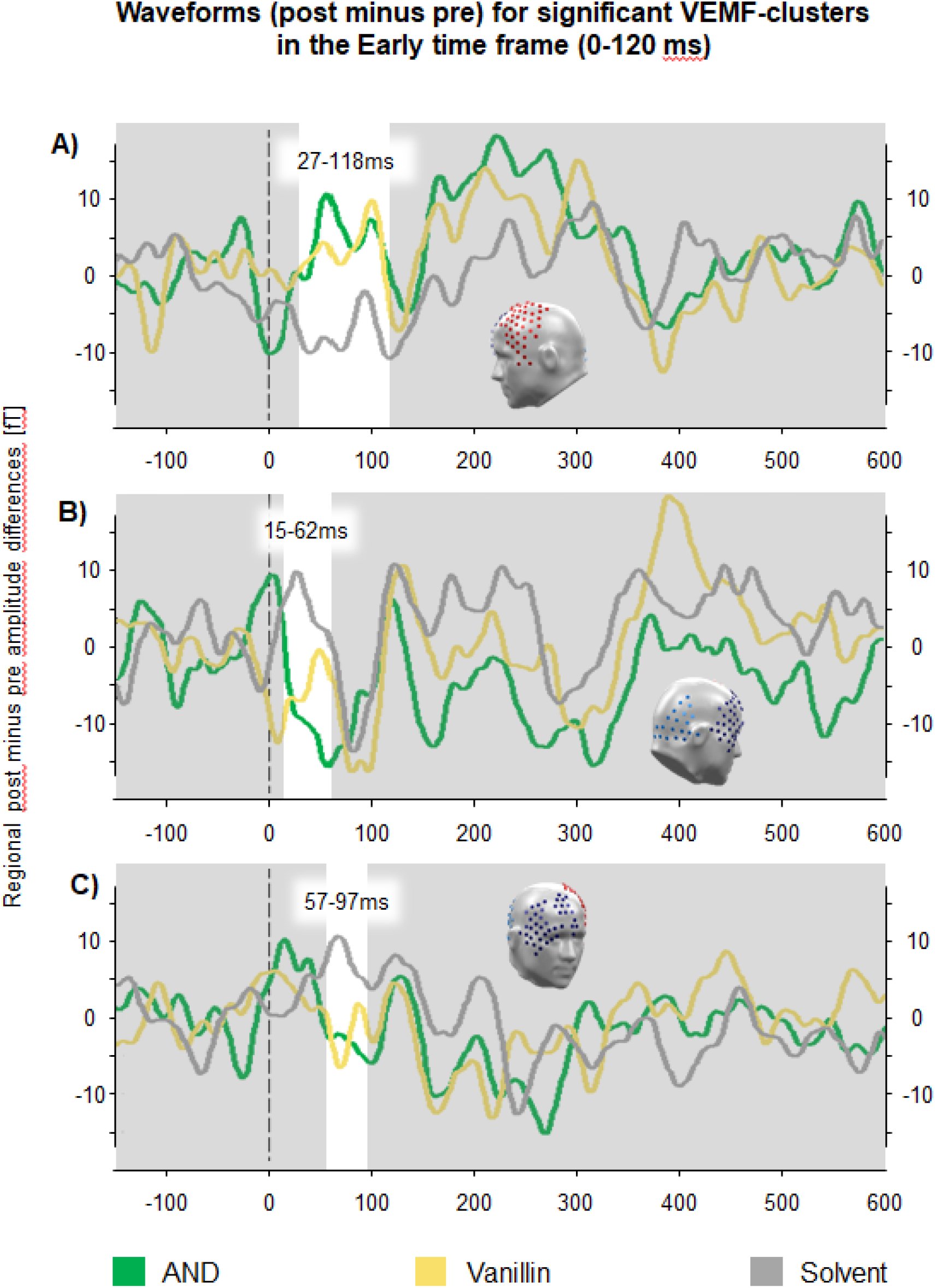
Waveforms for the significant VEMF clusters in (A) 27-118 ms, (B) 15-62 ms, and (C) 57-97 ms within the early (0-120 ms) time frame. Panels illustrate the regional post minus pre amplitude differences in femtotesla (fT) in a time frame of 150 ms before stimulus onset (baseline) to 600 ms after stimulus onset. Significant time frames are illustrated by a white background (after correction for multiple comparisons).

#### 4.2.2 Early posterior negativity (EPN-m)

In the mid-latent time frame (120-300 ms), a left hemispheric (143-230 ms) cluster and a subsequent but temporally overlapping right hemispheric cluster (202-263 ms), reached significance (CCM: 1758; left CM: 6796.3; right CM: 3577.9; Fig. 3B and Fig. 5). As in the early interval, the left cluster showed increased outgoing fields (positivity) for both AND- and VAN-associated faces, compared to SOLV-associated faces (AND vs. SOLV *t*(19) = 3.85, *p* = .001; VAN vs. SOLV *t*(19) = 3.49, *p* = .002; Fig. 3B, left; Fig. 5A), while the right cluster showed increased ingoing fields (negativity) for this comparison (AND vs. SOLV *t*(19) = −5.59, *p* < .0001; VAN vs. SOLV *t*(19) = −4.4, *p* < .0001; Fig. 3B, right; Fig. 5B). Again, AND- and VAN-associated faces did not differ, neither in the left (AND vs. VAN *t*(19) = −.1, *p* = .921) nor the right hemispheric cluster (AND vs. VAN *t*(19) = −.96, *p* = .347). We interpreted these components as an EPN-m, occurring in the typical EPN time window and yielding the classical difference topography.

**Figure 5:**
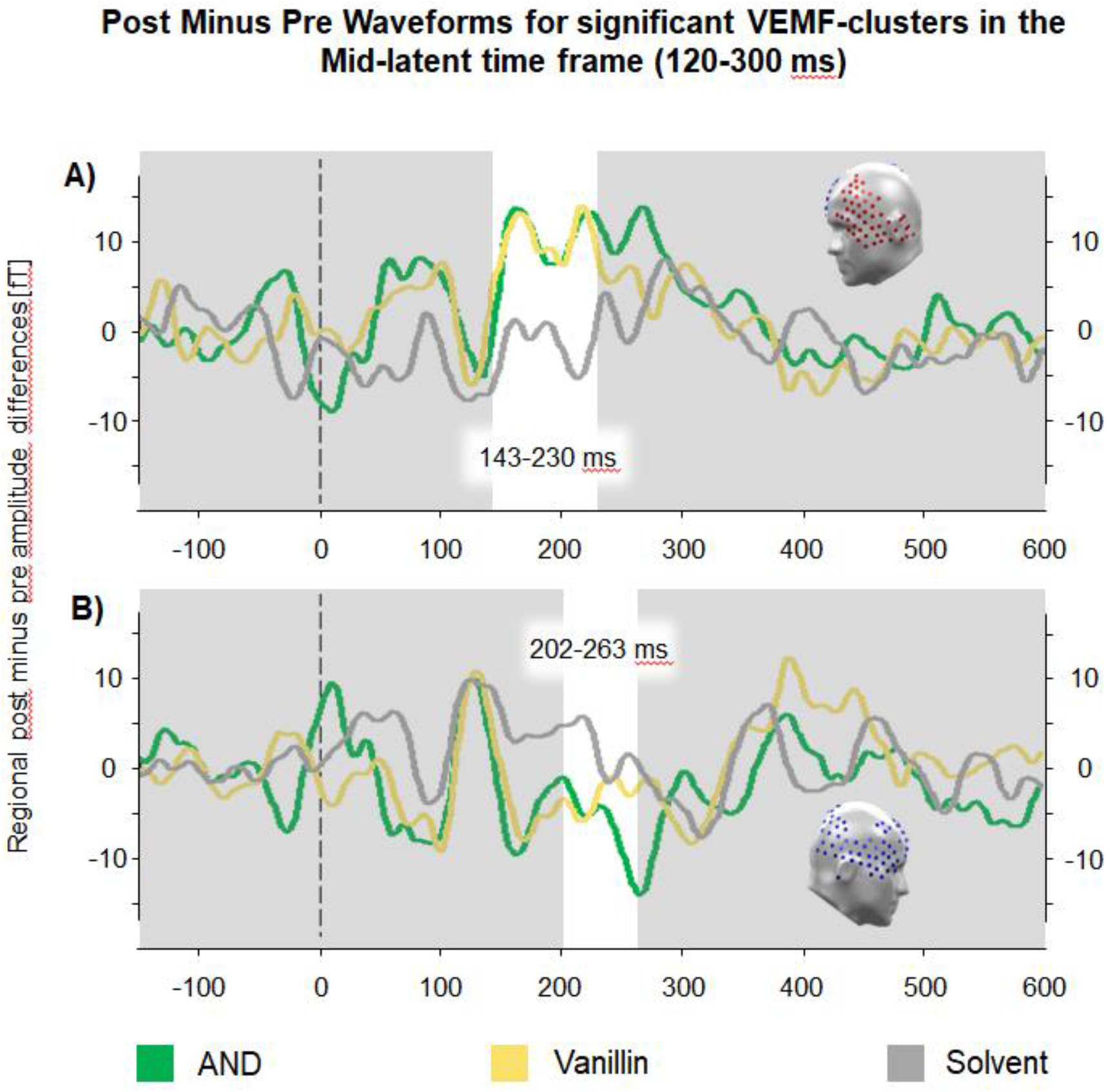
Analogous to Fig. 4, panels illustrate waveforms for the significant VEMF clusters in (A) 143-230 ms and (B) 202-263 within the mid-latent (120-300 ms) time frame.

## 5 Discussion

We investigated whether AND, a metabolite of testosterone found mainly in male sweat and consciously perceived as neutral, acts as an effective social chemosignal for human mating behavior in young female participants. We used AND as an unconditioned stimulus in a male face-learning paradigm and assessed neural and behavioral changes in participants’ perception of AND-associated faces compared to faces conditioned with a consciously positive odor (VAN) and with a neutral control odor (SOLV). If AND acts as an effective social chemosignal, it should evoke typical emotional and attentional learning effects that are qualitatively comparable to the pleasant stimulus VAN.

To the best of our knowledge, this is the first study demonstrating neural as well as behavioral effects of AND in a learning paradigm, potentially indicating its function as a social (mating) chemosignal with long-lasting effects. As expected, typical correlates of enhanced emotional attention for AND- and VAN-associated faces were found in early (15-118 ms) and mid-latent intervals (143-263 ms). Importantly, comparably amplified neural effects were observed for both AND and VAN compared to SOLV, despite participants having rating AND as a neutral odor (at the same level as SOLV), and despite participants’ lack of contingency awareness of CS-US pairings. Converging with the neural effects, AND- and VAN-paired faces achieved higher valence ratings in the post-conditioning face ratings than did the SOLV-paired faces. We will discuss these effects in more detail in the following.

### 5.1 Neural effects

In the early time frame, three components were observed from 15 to 118 ms. The temporal and spatial distribution as well as the polarity of these effects conform to components that indicate rapid salience detection and emotional attention (Hintze et al., 2014; Steinberg et al., 2012; Stolarova et al., 2006). Our data, therefore, suggest early enhanced emotional attention to faces paired with AND and VAN compared to faces paired with SOLV alone. While early correlates of enhanced emotional attention to conditioned faces has already been demonstrated for consciously aversive UCS odors without CS-US contingency awareness (Steinberg et al., 2012) as well as for consciously aversive and pleasant US odors with CS-US contingency awareness (Steinberg et al., 2013a), our results replicate and extend these findings toward consciously appetitive odors without CS-US contingency (here VAN). For faces paired with the perceptually neutral AND, our findings underscore the idea that AND seems to convey affective information, presumably of a social nature with a potential impact on mating behavior.

The two components evident in the mid-latent time frame reveal a timing and a topography corresponding to the predicted EPN-m (143-263 ms; positivity for AND- and VAN-associated faces, compared to SOLV-associated faces over the left hemisphere, and a relative negativity over the right hemisphere). As the EPN-m is a typical correlate of emotional attention (Schupp et al., 2006), these results support the interpretation that AND may carry emotional social information. Additionally, an estimation of neural sources (see supplemental information; Table S1) revealed spatially and temporally distributed clusters within seven regions, further supporting the assumption of preferred processing of AND- and VAN-associated faces compared to SOLV-associated faces^4^.

### 5.2 Behavioral and self-report effects

Neural effects might also be explained by mere physiological arousal, which AND might mediate. However, this interpretation contradicts face-rating results showing that arousal scores were not modulated by CS-US associations, as faces previously paired with either AND or VAN were rated as more appetitive than faces associated with SOLV alone. For VAN-paired faces, these results further replicate and extend our previous finding regarding aversive olfactory MultiCS conditioning, which showed that faces paired with a consciously aversive odor, without CS-US contingency awareness, were rated as less appetitive than faces associated with a neutral odor (Steinberg et al., 2012). Importantly, the higher valence scores for AND-paired faces are independent of the odor’s perceptual valence, as participants rated AND (and SOLV) as being a neutral odor, whereas they rated VAN as being appetitive. Our interpretation is that behavioral as well as neural effects occur independent of the perceptual odor valence but are modulated by an emotional content carried by AND, which seems to exert early effects possibly relevant for mating behavior.

In addition to strong support for AND as a social chemosignal, these findings also provide important information with respect to mechanisms of affective associative learning. While previous MultiCS studies revealed that awareness of CS-US contingency is not a necessary precondition for successful associative learning, the current results demonstrate that explicit perception of the US is also not a precondition (Bröckelmann et al., 2013, 2011; Junghöfer et al., 2017, 2015; Rehbein et al., 2015, 2014; Steinberg et al., 2013a, 2013b, 2012).

### 5.3 Limitations

When asked to rate the odor’s intensity, participants reported AND to be as intense as VAN, and both to be more intense than SOLV alone. One might thus question whether neural effects could be explained by mere intensity differences between the odors. It might be possible (but, as far as we know, not yet investigated) that pairing a CS with a more intense but otherwise equally neutral US could result in a relatively enhanced CS-US association, which would be at least qualitatively similar to learning effects driven by aversive or appetitive US relative to neutral US. However, such hypothetical US intensity-driven learning effects should be significantly smaller than US valence-driven ones, but our study revealed behavioral and neural effects of comparable strength for AND and VAN relative to SOLV alone. Moreover, post hoc linear regressions (see supplemental information; Table S2 and Table S3) revealed estimated neural activity differences within all significant spatio-temporal clusters to be independent of individual US intensity ratings. Participants could not indicate face-odor pairings above chance level, showing that CS-US learning occurred despite a lack of contingency awareness (Ohman and Mineka, 2001; Rehbein et al., 2014; Steinberg et al., 2012).

Given our hypothesis-driven approach that predicted effects in heterosexual women, we restricted our investigation to this group. Future research could next investigate the interesting question of whether and how these social learning mechanisms depend on sex and sexual orientation.

There is mounting evidence that high levels of testosterone are associated with “the search for, and maintenance of, social status” in humans (Eisenegger et al., 2011). Since AND is a metabolite of testosterone, the enhanced processing of AND-associated faces might also reflect that stronger emotional attention is allocated to individuals with an assumed higher social status. However, social status of males is also an important attractiveness factor for women (Sadalla et al., 1987). Again, future studies with heterosexual male and heterosexual female participants might help to resolve whether AND contains information specifically relevant for human mating, beyond its impact of social status.

As we did not control for participants’ phase in their menstrual cycle, we cannot say anything about potentially stronger effects in the most fertile phase. To reduce potential variance, we restricted our study to females not using hormonal contraception. It would, of course, be interesting to investigate whether hormonal contraception modulates effects of AND.

The concentration of AND in our study was 46 ppm, which is up to 170 pmol per mg of SOLV. Nixon and colleagues (1988) found that the AND concentrations on axillary hair of forty males was between 0 and 142 pmol/mg, with strong variability among individuals. However, our study and the study by Nixon and colleagues (1988) did not measure the relevant amount of AND molecules in the breathing air of the potential chemical signal receiver. More specific measures of AND concentration in the proximity of males would be needed to decide whether our AND concentration conforms to a natural physiological range and to investigate whether there is a maximal distance between chemosignal emitter and receiver for effective associative learning (are physical closeness and/or nakedness necessary conditions, or does AND affect even everyday social communication?).

Next, research on human chemosignals has primarily focused on AND, but it might be worthwhile to assess whether other components of male sweat or male sweat in itself induce similar effects in women.

As a first approach, we used social CS (faces), thus providing an AND association with high ecological validity. We do not know whether other non-social stimuli (e.g., Garbor gratings) could also be associated with AND, but a successful association with non-social CS would not undermine the interpretation of AND as a social chemosignal.

A N of 20 participants may be considered as a small sample size. However, the sample of participants was well controlled including only women using no hormonal contraception. Even more importantly, we employed a specifically developed experimental design using a within-subject design in a MultiCS conditioning paradigm. This allows for an excellent signal-to-noise ratio and is thus perfectly suited for recording ERPs or EMRFs (Bröckelmann et al., 2011; Steinberg et al., 2013b, 2012). The strong electrophysiological effects found in this study confirm the validity of this approach. Moreover, a follow-up study of our lab using human chemosignals of anxiety instead of AND in the same paradigm replicated the findings of this study.

## 6 Conclusions

We present data showing that the testosterone metabolite AND, which is consciously perceived as a neutral odor, modulates motivated attention and attractiveness ratings of AND-associated males and, thus, might influence human mating behavior. AND evoked learning effects as strongly as the pleasant stimulus VAN according to neural and behavioral measures. Despite participants’ lack of contingency awareness of CS-US pairings, they rated faces paired with AND or VAN as more pleasant than SOLV-paired faces. In other words, AND evoked neural and behavioral learning effects as strongly as an unconditioned pleasant stimulus, even though AND was not consciously perceived as being pleasant, but neutral. These findings support the interpretation that AND is, in fact, an effective human social chemosignal.

## Supporting information

Supplemental data items

## 8 Acknowledgements

This work was funded by the Deutsche Forschungsgemeinschaft grant SFB TRR-58 C01. For the many productive discussions, we want to thank Ludger Elling, Christian Steinberg, Maimu Rehbein, and Ida Wessing. Thanks also to Andreas Wollbrink, Mikheil Gogiashvili, Karin Berning, Hildegard Deitermann, and Ute Trompeter for their technical assistance and help with the data collection. Thanks also to Brian Bloch and Celeste Brennecka from the Science Writing Support Service from the University of Münster for proofreading. Portions of the research in this paper use the FERET database of facial images collected under the FERET program, sponsored by the DOD Counterdrug Technology Development Program Office. The authors would also like to thank Daniel Lundqist, Anders Flykt, and Arne Öhman for their permission to use the Karolinska Directed Emotional Faces (KDEF) database. Development of the MacBrain Face Stimulus Set / NimStim set of facial expressions was overseen by Nim Tottenham and supported by the John D. and Catherine T. MacArthur Foundation Research Network on Early Experience and Brain Development. Please contact Nim Tottenham at for more information concerning the stimulus set.

## Footnotes

1 Although the odor stimulation device was positioned outside the shielded room, its pressure valves were perceivable as soft sounds inside the chamber. To prevent even slight additional audio-visual associative leaning, participants were presented with a continuous acoustic white noise that masked these sounds. At the start, participants were asked to tune the noise volume to be as loud as possible without it being annoying.

2 As participants were typically exhaling in the second half of CS presentation, US presentation was restricted to the first half, facilitating the subsequent odor neutralization.

3 As there were no specific predictions regarding the relative effect strength of AND and VAN conditioning (see introduction), both were weighted equally.

4 Effects of associative learning in the so-called source space (i.e., estimated neural sources based on inverse modeling) generally support the findings in sensor space but are more manifold and complex than the quite unambiguous sensor-space findings. As identifying underlying structures was not our aim, we decided to focus on the sensor space effects for the sake of comprehensibility, and we report the source space findings as supplemental information (Table S1).

