## Supplemental data items for "Women show enhanced attractiveness ratings and neural processing of male faces associated with male chemosensory signals"

*Table S1:* Results of critical cluster masses (CCM), cluster masses (CM) and paired *t*-tests for significant ($p$<.05 sensor level; $p$<.01 cluster level) MNE clusters (estimated neural sources). Related to Figure 3

| CCM | CM | AND vs VAN | AND vs SOLV | VAN vs SOLV |
|---|---|---|---|---|
| | right TPJ (37-82ms): AND < SOLV & VAN < SOLV | | | |
| | 645.5 | $t(19) = 2.25$ $p = .037$ | $t(19) = -3.87$ $p = .001$ | $t(19) = -7.05$ $p < .0001$ |
| | right occipito-parietal (22-40ms): AND > SOLV & VAN > SOLV | | | |
| | 385.1 | $t(19) = 0.51$ $p = .614$ | $t(19) = 4.38$ $p = .001$ | $t(19) = 3$ $p = .007$ |
| 137.5 | occipital (5-35ms): AND > SOLV & VAN > SOLV | | | |
| | 202.5 | $t(19) = -.12$ $p = .907$ | $t(19) = 4.56$ $p < .0001$ | $t(19) = 3.73$ $p = .001$ |
| | potentially right fusiform gyrus (10-30ms): AND > SOLV & VAN > SOLV | | | |
| | 155.3 | $t(19) = .95$ $p = .353$ | $t(19) = 3.52$ $p = .002$ | $t(19) = 3.7$ $p = .002$ |
| | left dlPFC (275-300ms): AND > SOLV & VAN > SOLV | | | |
| | 824.7 | $t(19) = -.67$ $p = .512$ | $t(19) = 3.89$ $p = .001$ | $t(19) = 5.36$ $p < .0001$ |
| 288.5 | right medial temporal cortex (120-147ms): AND > SOLV & VAN > SOLV | | | |
| | 453.5 | $t(19) = -.77$ $p = .454$ | $t(19) = 5.73$ $p < .0001$ | $t(19) = 5.92$ $p < .0001$ |
| | vmPFC (223-242ms): AND > SOLV & VAN > SOLV | | | |
| | 306.3 | $t(19) = -.61$ $p = .548$ | $t(19) = 3.06$ $p = .006$ | $t(19) = 3.14$ $p = .005$ |

*Table S2:* Results of linear regression of odor intensity rating and neural activation in early VEMF clusters (0-120ms) show no influence of odor intensity on neural activation. Related to Figure 2 and 3

| | B (SE) | Lower 95 % CI | Upper 95 % CI |
|---|---|---|---|
| left frontal cluster (27-118ms) | | | |
| Intercept | -4.19(6.67) | -17.11 | 8.94 |
| Intensity | -0.19(0.15) | -0.49 | 0.11 |
| Activity AND | 12.66(12.90) | -13.46 | 38.30 |
| Activity VAN | -13.40(16.05) | -44.68 | 18.65 |
| Intensity X Activity AND | 0.16(0.22) | -0.27 | 0.60 |
| Intensity X Activity VAN | 0.51(0.26) | 0.02 | 1.00 |
| right posterior cluster (15-62ms) | | | |
| Intercept | -0.59(6.30) | -13.04 | 11.44 |
| Intensity | 0.21(0.15) | -0.08 | 0.52 |
| Activity AND | 11.16(12.45) | -13.47 | 34.74 |
| Activity VAN | -20.87(15.23) | -51.34 | 8.23 |
| Intensity X Activity AND | -0.52(0.22) | -0.93 | -0.08 |
| Intensity X Activity VAN | 0.00(0.25) | -0.47 | 0.49 |
| right frontal cluster (57-97ms) | | | |
| Intercept | 8.07(6.40) | -4.24 | 20.84 |
| Intensity | 0.06(0.15) | -0.24 | 0.34 |
| Activity AND | -19.46(12.37) | -44.33 | 4.53 |
| Activity VAN | -12.49(15.64) | -45.29 | 17.67 |
| Intensity X Activity AND | 0.06(0.22) | -0.36 | 0.48 |
| Intensity X Activity VAN | -0.07(0.25) | -0.53 | 0.44 |

*Table S3:* Results of linear regression of odor intensity rating and neural activation in mid-latent VEMF clusters (120-300ms) show no influence of odor intensity on neural activation. Related to Figure 2 and 3

| | B (SE) | Lower 95 % CI | Upper 95 % CI |
|---|---|---|---|
| left cluster (143-230ms) | | | |
| Intercept | -4.19(6.67) | -17.11 | 8.94 |
| Intensity | -0.19(0.15) | -0.49 | 0.11 |
| Activity AND | 12.66(12.90) | -13.46 | 38.30 |
| Activity VAN | -13.40(16.05) | -44.68 | 18.65 |
| Intensity X Activity AND | 0.16(0.22) | -0.27 | 0.60 |
| Intensity X Activity VAN | 0.51(0.26) | 0.02 | 1.00 |
| right cluster (202-263ms) | | | |
| Intercept | 8.07(6.40) | -4.24 | 20.84 |
| Intensity | 0.06(0.15) | -0.24 | 0.34 |
| Activity AND | -19.46(12.37) | -44.33 | 4.53 |
| Activity VAN | -12.49(15.64) | -45.29 | 17.67 |
| Intensity X Activity AND | 0.06(0.22) | -0.36 | 0.48 |
| Intensity X Activity VAN | -0.07(0.25) | -0.53 | 0.44 |

### Supplemental Experimental Procedures

#### *Methods - Participants*

Participants were asked, by means of an in-house screening questionnaire, to report their sexual orientation and handedness. They were also asked to report all medication they were taking. Such medication was then checked for possible effects on the sense of smell.

#### *Analysis - Estimation of neural sources*

Averaged responses were used to calculate the L2-Minimum-Norm-Estimates (L2-MNE) method (Hämäläinen and Ilmoniemi, 1994) for cortical sources. The L2-MNE is an inverse modeling technique, enabling the estimation of distributed neuronal network activity, without the need for a priori specifications of the location and/or number of active current dipoles (Hauk, 2004). As the source model, a spherical shell with evenly distributed 2 (azimuthal and polar direction) x 350 dipoles was used, with a source shell radius of 87% of the individually fitted head radius, which approximately corresponds to the gray matter depth. Across all participants and conditions, a Tikhonov regularization parameter *k* of 0.1 was used. Analogous to the analysis of the VEMFs, baseline measurement MNE data were subtracted from extinction measurement MNE data and weighted with contrasts +1 (AND), +1 (vanillin) and -2 (solvent). For the selection of significant clusters, the same criteria and time frames were applied as in the VEMF data.
